# OmniDetect: a plant-produced universal IgG binder replacing secondary antibodies in immunoassays

**DOI:** 10.64898/2026.01.14.699418

**Authors:** Aditya Prakash Soni, Madhu Kumari, Thangarasu Muthamilselvan, Youra Hwang, Seungjin Woo, Inhwan Hwang

## Abstract

High sensitivity in immunodetection technologies has long been a priority for both diagnostic kit development and routine laboratory research. We developed a recombinant fusion protein, OmniDetect-G3, as an alternative to conventional HRP-conjugated secondary antibodies, and successfully produced it in plants. OmniDetect-G3 binds to primary IgG antibodies regardless of host species, and also recognizes chicken IgY. In comparative analyses, OmniDetect-G3 exhibits 165–381% higher sensitivity than commercial secondary antibodies in western blot analysis and 30%-1145% higher sensitivity in ELISA, depending on commercial products tested. Moreover, OmniDetect-G3 showed robust performance when used with sera from vaccinated animals (porcine and bovine). The chemiluminescence signal generated by OmniDetect-G3 remained stable up to 30 minutes after reaction initiation. These findings indicate that plant-produced OmniDetect-G3 has strong potential as a universal mimetic to animal-derived HRP-conjugated secondary antibodies, offering simplified immunoassay workflows and improved detection efficiency.

## Introduction

There has been extensive effort to improve immunodetection systems in both laboratory-based scientific research and commercial diagnostic kits^1^. Higher detection sensitivity creates new opportunities in the diagnostic application^2^, particularly when target antigens or antibodies are present at extremely low levels, making accurate detection challenging^3^. Among the most widely used detection approaches is the use of HRP-conjugated secondary antibodies. However, these reagents have several limitations ^4^, including suboptimal sensitivity, the requirement for species-matched specificity to primary antibodies, and batch-to-batch variation in performance. In addition, production of these secondary antibodies requires multiple steps-including purification of HRP, purification of anti-IgG antibodies, and cross-linking of HRP and anti-IgG antibodies, followed by purification of the conjugated products-which collectively increase manufacturing costs. Another significant concern is the reliance on animals for the production of anti-IgG antibodies. Given that the global antibody market exceeds USD 1 billion annually (https://tinyurl.com/mywm2as6), the number of animals used for generating secondary antibodies is likely substantial.

As an animal-free alternative, nanobody-based HRP-conjugated proteins have been introduced for similar application purposes ^5–7^. Antigen-specific nanobodies can be generated either through immunization of animals ^8^ or via an in vitro screening approach^9^. These nanobodies can then be genetically fused to HRP, allowing nanobody-HRP fusion proteins to be produced in heterologous expression systems. However, nanobody-based secondary antibodies have intrinsic limitations. In general, nanobodies exhibit lower binding affinity to their targets compared to IgG antibodies, and each nanobody is specific to a particular IgG. This restricts the development of nanobody-based secondary antibodies^10–12^.

Another candidate for universal IgG detection is protein A^13,14^ or protein G^15,16^, both of which bind to IgG molecules from a wide range of animal species. Notably, a recent study by Song et.al (2022)^17^ demonstrated that a GB1-fused recombinant protein produced in plants can be directly detected by various HRP-conjugated secondary antibodies that are typically specific for rat, rabbit, or mouse IgG in western blot analyses. Earlier studies have similarly exploited the IgG-binding property of GB1 to develop universal IgG binders by several laboratories^18–20^. However, in most cases, the sensitivity of these GB1-based proteins did not match that of commercial secondary antibodies^21^. We also chose GB1 and HRP as key components for developing a universal binder capable of recognizing IgG from various animal species. However, based on the earlier studies, a major challenge we need to address is the inherently weaker binding affinity of GB1 to IgG compared with that of conventional anti-IgG antibodies.

In this study, we compensated for the weaker intrinsic binding affinity of GB1 to IgG by introducing avidity-based enhancements at two levels. First, we fused multi-copy GB1 repeats to HRP, and second, we enabled multimerization of the fusion protein into a higher-order complex. Through this strategy, we successfully developed novel composite proteins-most notably *OmniDetect-G3* (Universal Detection), which exhibit significantly higher sensitivity toward various IgG antibodies compared to commercial secondary antibodies. OmniDetect-G3 also binds to IgY, though with a somewhat lesser degree of sensitivity than IgY-specific secondary antibody. In addition, we achieved high-level production in plants. Collectively, *OmniDetect-G3* overcomes long-standing limitations in HRP-conjugated secondary antibodies-including host-species dependency, reliance on animal immunization, and batch-to-batch variation-thereby representing both a conceptual and technical breakthrough in immunodetection.

## Results

### Construction of OmniDetect-G1 and validation of the hypothesis

For developing an agent capable of detecting IgGs in an immunoassay, we designed a fusion protein consisting of the GB1 domain and HRP. However, the binding affinity of the GB1 domain to IgGs is in the micromolar range, whereas IgG-specific antibodies typically bind in the nanomolar or sub-nanomolar range IgGs^22^. These weaker affinities result in reduced sensitivity, and previous studies have attempted to compensate for this by using multiple copies of GB1 through various approaches^23^. Similarly, we employed the multiple copies of the GB1 domain to overcome its lower binding affinity through a two-layer avidity strategy: first, by fusing multiple GB1 repeats with HRP, a single molecule, and second, by inducing multimerization of the fusion protein. Multimerization of a fusion protein consisting of multiple GB1 domains and an HRP moiety results in a complex with a greatly increased valency of GB1 and multiple HRP enzymes, potentially enhancing both IgG binding strength and signal intensity. To test this concept, we examined multimerization domains such as Foldon^24^ (forms trimer), PilZ^25^ (forms tetramer), and CTB^26^ (forms pentamer). We constructed a CTB-based complex from a fusion containing 3 copies of GB1 and one copy of HRP termed OmniDetect-G1 (Fig. 1a). The GB1 used was an ERH mutant known to exhibit higher binding affinity to human IgG (nanomolar *Kd*)^27^. In this design, the assembled OmniDetect-G1 complex is expected to contain 15 copies of GB1 and 5 copies of HRP, conferring much higher sensitivity for IgG detection compared with a monomeric fusion protein.

**Fig. 1.**
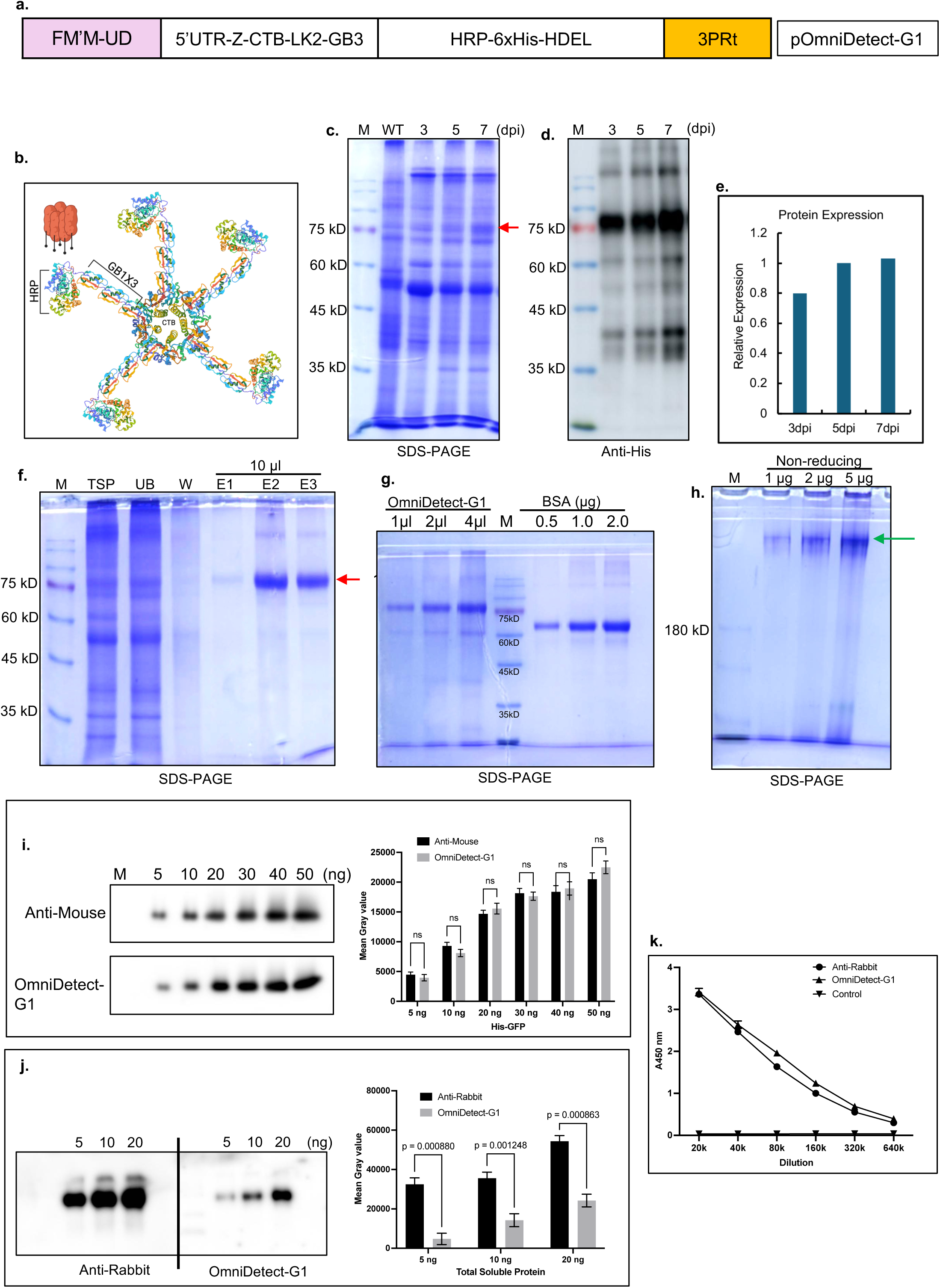
Initial design and validation of the first OmniDetect hypothesis. **a.** Schematic representation of the construct, *pOmniDetect-G1*. FM’M-UD, an artificial promoter; Z, γ-zein signal peptide (1-19 aa, Q548E8); CTB, cholera toxin-B subunit (WP_338286491.1); LK2, linker GGGGSx2; GB3, three copies of B1 domain of Streptococcal protein G; HRP, horseradish peroxidase; 6xHis, His-tag; HDEL, ER-retention signal; 3PRt, artificial composite terminator. **b.** Conceptual design of OmniDetect-G1 generated in BioRender using structures imported from the PDB database. **c-e.** Expression of Omnidetect-G1. Leaf samples were collected at 3, 5, and 7 dpi; total soluble protein was extracted, separated on 10% SDS-PAGE, and analyzed by CBB-staining and western blot using Anti-His IgG. Relative expression levels were quantified as mean gray values of the target band and plotted. **f-h.** Purification, quantification, and multimer analysis of OmniDetetc-G1 protein. M, protein marker; TSP; total soluble protein; UB, unbound fraction; W washing fraction; E, elution fractions (E1, 50 mM imidazole; E2, 100 mM; E3, 250 mM). The arrow indicates the target protein band. **i-j.** Side-by-side comparison of commercial HRP-conjugated anti-mouse and anti-rabbit secondary antibodies with OmniDetect-G1 in a western blot. **k.** ELISA comparing commercial anti-rabbit secondary antibodies with OmniDetect-G1. Data are represented as mean ± SE (n = 3). Error bars indicate standard error. P-values were calculated by Student’s t-test. The OmniDetect-G1 conceptual protein structure was created using Biorender.com. PDB ID: CTB: 1PZJ, GB1: 1GB1, HRP: 1HCH.

OmniDetect-G1 was successfully expressed in *Nicotiana benthamiana* leaves using *Agrobacterium*-mediated transient expression with peak accumulation at 5 dpi (Fig. 1c-e). The target protein band appeared at approximately ∼80 kDa,-slightly higher than the predicted size (∼75 kDa), likely due to glycosylation^28^. HRP is known to be heavily glycosylated, which is important for its enzymatic activity^29^. OmniDetect-G1 was purified by Ni^2+-^NTA chromatography (Fig. 1f) and quantified (Fig. 1g) using serially diluted BSA standards. Complex formation was confirmed under non-reducing conditions (Fig. 1h). Hemin treatment^30^ further enhanced HRP activity (Supplementary Fig. 1).

Next, before proceeding with the validation of OmniDetect-G1 by immunodetection assays, we compared various commercially available HRP-conjugated secondary antibodies and picked the best one for further experiments (Supplementary Fig. 2). We compared the IgG detection performance of OmniDetect-G1 with commercial HRP-conjugated anti-rabbit and anti-mouse IgG antibodies in both Western blotting and ELISA. In Western blot analysis, OmniDetect-G1 exhibited sensitivity comparable to that of HRP-conjugated anti-mouse IgG antibodies in the 5 - 50 ng range of target proteins, but its sensitivity was 50% - 80% lower than that of HRP-conjugated anti-rabbit IgG antibodies, depending on the target protein amount (Fig. 1i-j). Interestingly, in contrast to western blot results, OmniDetect-G1 generated higher signal intensity than the commercial HRP-conjugated anti-rabbit IgG by 2 to 31% depending on dilution (1:20,000 – 1:640,000), in ELISA (Fig. 1k). At present, the reason for the discrepancy between western blot analysis and ELISA performance is unclear. One possibility is that the accessibility of OmniDetect-G1 to the primary antibodies differs between the two assay formats.

### Replacing the CTB with Tenascin-C N-terminal domain enhances signal strength in both western blot analysis and ELISA

We aimed to improve the detection sensitivity of OmniDetect so that it would become comparable to commercially available secondary antibodies. One approach was to increase the degree of multimerization of the GB1-HRP fusion protein. Another consideration was the flexibility of HRP; in OmniDetect-G1, the GB1 repeats and HRP are likely positioned very close to each other due to the structural conformation of CTB. A previous study suggested that when the HRP moiety is present in excess during western blot imaging, it may quench chemiluminescent reactions^31^, thereby reducing signal intensity. To address this issue, we selected Tenascin-C, a large ECM glycoprotein^32^, in which two trimers assemble to form a hexamer ^33^. The hexameric structure has radiating C-terminal arms that radiate outward. The N-terminal domain of Tenascin-C (34-139 aa), referred to as TN106, is sufficient for hexamer formation^34^.

We designed OmniDetect-G2 (Fig. 2a-b) by replacing CTB in OmniDetect-G1 with TN106 as the multimerization domain. OmniDetect-G2 was expressed in *N. benthamiana*. Protein expression level remained comparable up to 7 dpi (Fig. 2c-e). Therefore, plants were harvested at 3-4 dpi. A protein band corresponding to OmniDetect-G2 was observed at approximately ∼90 kDa, which is higher than the calculated molecular weight (∼75 kDa). OmniDetect-G2 was purified using Ni^2+^-NTA chromatography (Fig. 2f), quantified (Fig. 2g), and detected at higher molecular weight positions under the non-reducing condition (Fig. 2h), confirming multimer formation. Unlike OmiDetect-G1, this version showed no effect of hemin treatment (Supplementary Fig. 3), although the underlying reason remains unclear.

**Fig. 2.**
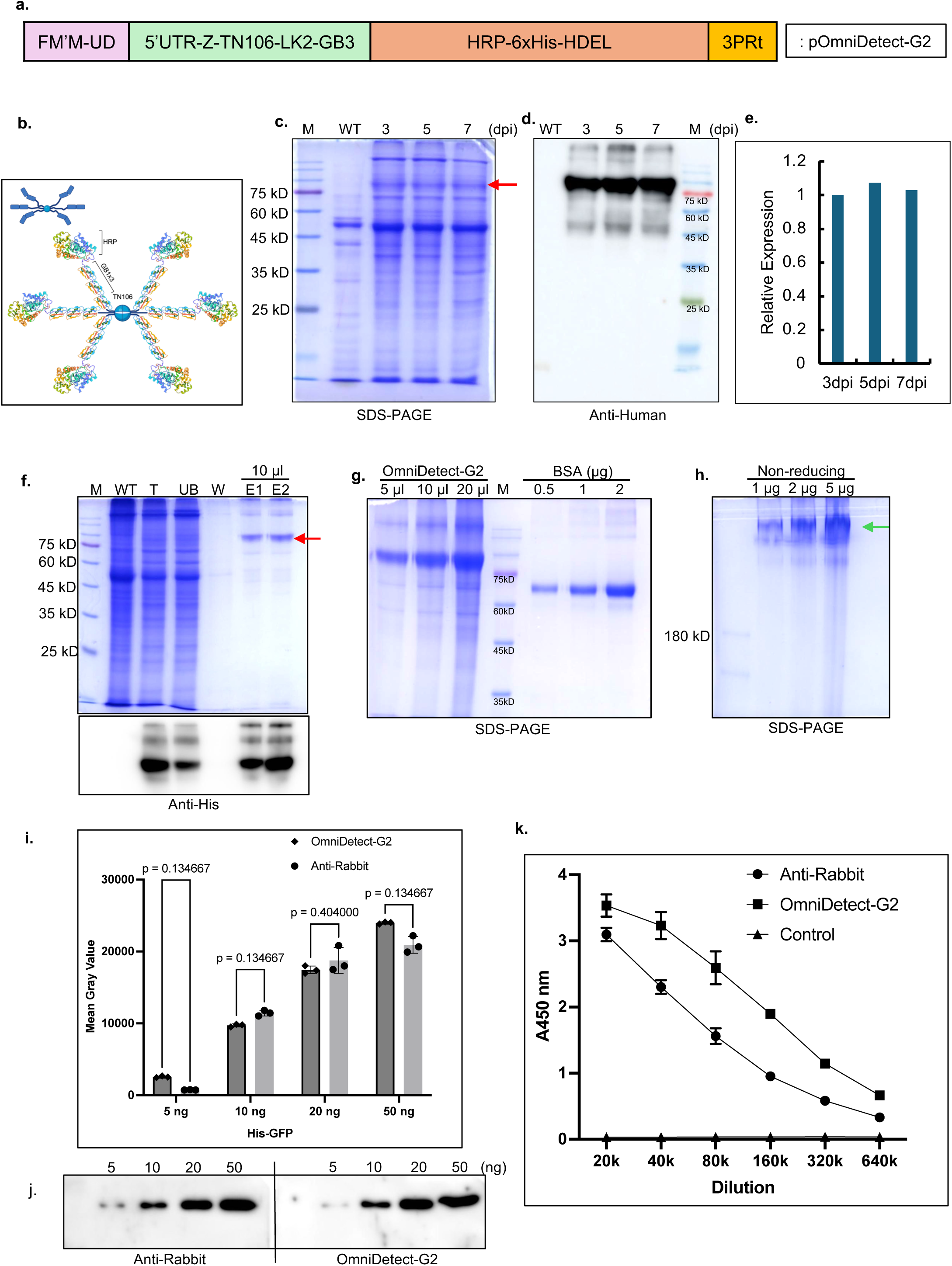
Construction and characterization of OmniDetect-G2. **a.** Schematic representation of the *pOmniDetect-G2* construct. FM’M-UD, an artificial promoter; Z, γ-zein signal peptide (1-19 aa, Q548E8); TN106, Tenascin-C N-terminal domain (34-139 aa, Uniprot. P10039); LK2, linker GGGGSx2; GB3, three copies of B1 domain of Streptococcal protein G; HRP, horseradish peroxidase; 6xHis, His-tag; HDEL, ER-retention signal; 3PRt, artificial composite terminator. **b.** Graphical representation of the Tenascin-C protein showing major regions, including the oligomer assembly domain. **c-e.** Expression of Omnidetect-G2. OmniDetect-G2 expression was analyzed as described in Fig. 1c-e. **f-h.** Purification, quantification, and multimer analysis of OmniDetect-G2 protein; **i-j.** Side-by-side comparison of commercial HRP-conjugated anti-rabbit secondary antibodies with OmniDetect-G2 in a western blot. Signal intensity was quantified as the mean gray values. **k.** ELISA comparing commercial anti-rabbit secondary antibody with OmniDetect-G2. Data represent mean ± SE (n = 3). Error bars indicate standard error. P-values were calculated by Student’s t-test. The conceptual OmniDetect-G2 protein structure was generated using BioRender.com. PDB ID: GB1: 1GB1, HRP: 1HCH.

Next, we evaluated the detection sensitivity of OmniDetect-G2 in western blot analysis. OmniDetect-G2 exhibited approximately 80-90% of the sensitivity of the commercial HRP-conjugated anti-rabbit IgG antibody when detecting 10 to 50 ng of target protein, and showed even higher sensitivity at 5 ng (Fig. 2i-j). However, overall, OmniDetect-G2 still did not match the performance of commercial secondary antibodies in Western blotting, and the results varied. In contrast, in ELISA, OmniDetect-G2 consistently outperformed the commercial HRP-conjugated anti-rabbit IgG antibody, demonstrating 14%-101% higher detection sensitivity, depending on the dilution factor (Fig. 2k).

To further improve the detection sensitivity, we tested mutant forms of HRP with previously reported higher enzymatic activities. Several HRP mutations were shown to enhance catalytic activity when expressed in *E. coli* and yeast. We generated two variants of OmniDetect-G2-G2-6m^35^ (6 mutations) and G2-4m^36^ (4 mutations)- by replacing the wild-type HRP with HRP6m and HRP4m, respectively (Supplementary Fig. 4a). Both variants were expressed in *N. benthamiana* and purified using Ni^2+^-NTA affinity chromatography (Supplementary Fig. 4b-e). OmniDetect-G2-6m and G2-4m exhibited approximately 70% (p<0.0001) lower sensitivity than OmniDetect-G2 in ELISA (Supplementary Fig. 4f). These results are consistent with the previous findings showing that plant-produced HRP with N-glycosylation possesses significantly higher enzymatic activity^37^ than HRP produced in *E. coli*.

### OmniDetect-G3, having one and two copies of GB1 on the N- and C-terminal of TN106, respectively, exhibits greatly enhanced detection sensitivity

We next aimed to improve the detection sensitivity of OmniDetect beyond that of commercial secondary antibodies. In OmniDetect-G1 and G2, three copies of GB1 were positioned in the middle of the fusion protein. We reasoned that these GB1 copies may experience steric restriction when binding to the Fc of the primary antibody, as they are flanked on both sides by large domains (CTB/TN106 and HRP. To address this, we relocated all three GB copies (GBx3) to the C-terminus of HRP (Supplementary Fig. 5a-d), followed by His-tag and the ER retention motif, HDEL. Several flexible GS-linkers were also inserted between HRP and GBx3 to increase domain mobility. Unexpectedly, this construct did not show any detectable signal in ELISA, suggesting that HRP in this fusion protein may not be active (Supplementary Fig. 5e). The cause of this inactivity of HRP remains unclear and was not further investigated.

Next, in another construct named OmniDetect-G3, we retained three copies of the GB1 domain, but rearranged so that one and two copies were positioned at the N and C-termini of TN106, respectively, followed by HRP at the C-terminus. This design was intended to provide the GB1 domains with better accessibility to the Fc region of the primary antibody. (Fig. 3a). When expressed in *N. benthamiana*, OmniDetect-G3 reached its highest expression at 3 dpi (Fig. 3b-d). OmniDetect-G3 was purified by using Ni^2+^-NTA affinity chromatography (Fig. 3e), quantified (Fig. 3f), and analyzed for multimer formation (Fig. 3g). We next examined the signal intensity of OmniDetect-G3 in both western blot analysis and ELISA. For western blotting, total soluble protein (expressing 6xHis-GFP) extracted from *N. benthamiana* was used. Total soluble proteins (50 and 10 ng) were separated by SDS-PAGE, subjected to western blot analysis. Membranes were probed with either rabbit anti-GFP or mouse anti-GFP antibodies as primary antibody, followed by incubation with commercial HRP-conjugated secondary antibodies or OmniDetect-G3. The dilution factor and concentration (1 mg/ml) were kept identical, and all imaging was performed simultaneously using a single auto-exposure setting. OmniDetect-G3 outperformed both commercial HRP-conjugated anti-rabbit and anti-mouse IgG antibodies in western blot analysis. The signal intensity produced by OmniDetect-G3 was 165-381% higher than that generated by the commercial HRP-conjugated secondary antibodies (Fig. 3h-i). Notably, OmniDetect-G3 showed markedly higher signal intensity when detecting lower amounts of target protein (10 ng of TSP), indicating that OmniDetect-G3 surpasses current commercial counterparts used in this study in detecting low-abundance proteins.

**Fig. 3.**
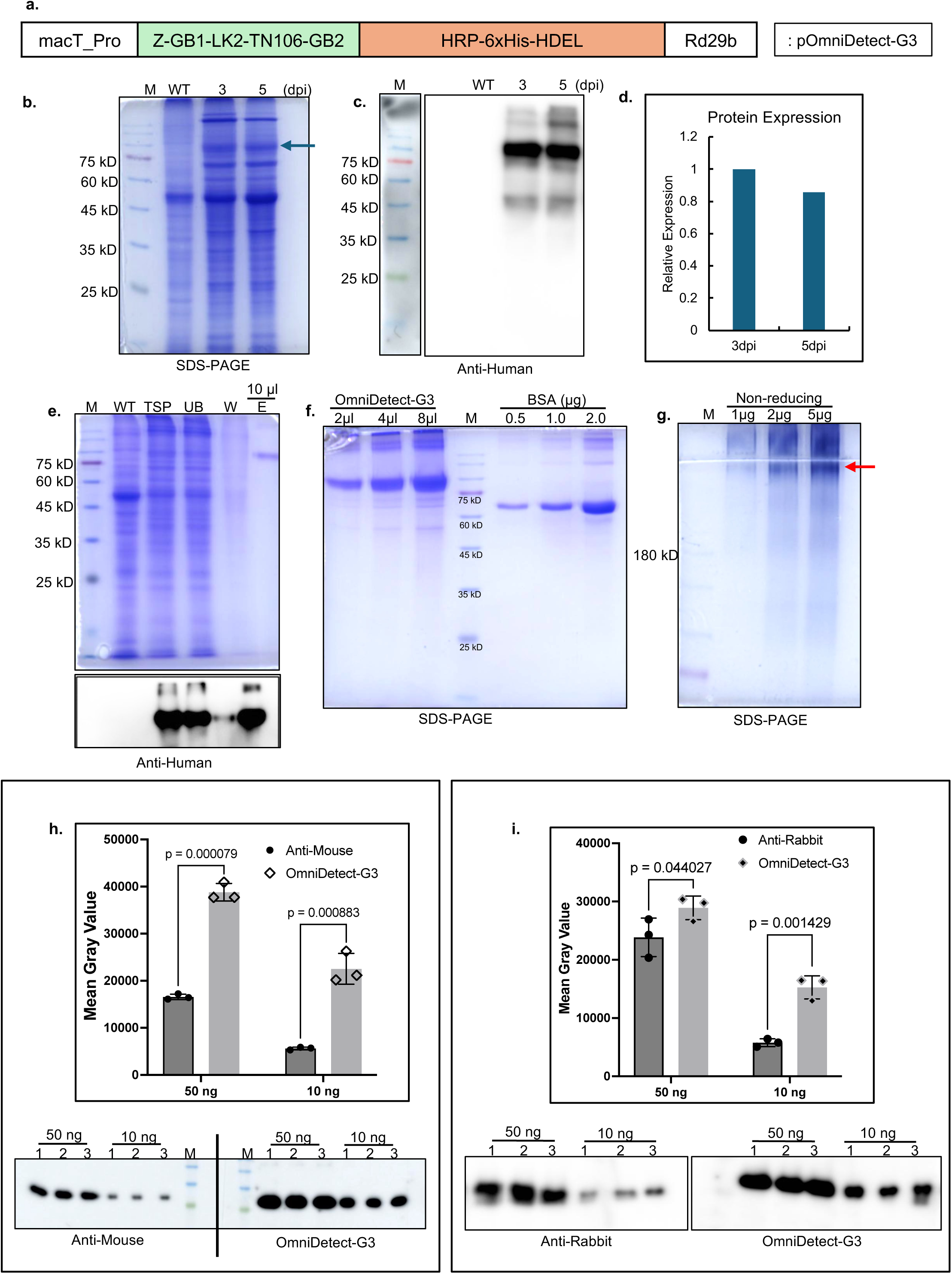
OmniDetect-G3 with one and two GB domains to the N and C-termini, respectively, of TN106 has a strongly enhanced signal in the detection of primary antibodies. **a.** Schematic representation of the construct, *pOmniDetect-G3*. macT, a chimeric promoter; Rd29B, *Arabidopsis RD29B* terminator; Z, γ-zein signal peptide (1-19 aa, Q548E8); GB1, single copy of B1 domain of Streptococcal protein G (GB1); TN106, Tenascin-c protein (34-139 aa, Uniprot. P10039); LK2, linker GGGGSx2; GB2, two copies of B1 domain of Streptococcal protein G; HRP, horseradish peroxidase; 6xHis, His-tag; HDEL, ER-retention signal. **b-d.** Expression of Omnidetect-G3. Leaf samples were collected at 3, 5, and 7 dpi; total soluble protein was extracted and separated on 10% SDS-PAGE, followed by CBB-staining and western blot using anti-human IgG. Relative expression was quantified as the mean gray values of the target band and plotted. **e-g.** Purification, quantification, and multimer analysis of OmniDetect-G3 protein. **h-i.** Side-by-side comparison of commercial HRP-conjugated anti-mouse and anti-rabbit secondary antibodies with OmniDetect-G2 in a western blot. Signal intensity was quantified as the mean gray values. Data represent mean ± SE (n = 3). Error bars indicate standard error. P-values were calculated by Student’s t-test.

### OmniDetect-G3 shows higher performance than commercial counterparts used in this study in detecting antibodies in ELISA

To further validate the versatility and sensitivity of OmniDetect-G3, we obtained rabbit, pig, and bovine sera containing anti-GFP, anti-CSF, and anti-tuberculosis Ag85A antibodies, respectively. These antibody levels were measured by ELISA using either commercial secondary antibodies (rabbit IgG, pig IgG, and bovine IgG) or OmniDetect-G3. In all three ELISAs, OmniDetect-G3 outperformed the counterparts used in this study to varying degrees. OmniDetect-G3 yielded 13%-30% higher signal intensity than HRP-conjugated anti-rabbit IgG (Fig. 4a), 475%-138% higher signal intensity than anti-pig IgG, and 52%-1145% higher signal intensity than commercial HRP-conjugated anti-bovine IgG antibody. These results confirm that OmniDetect-G3 possesses higher sensitivity than commercial secondary antibodies in both western blot analysis and ELISA.

**Fig. 4.**
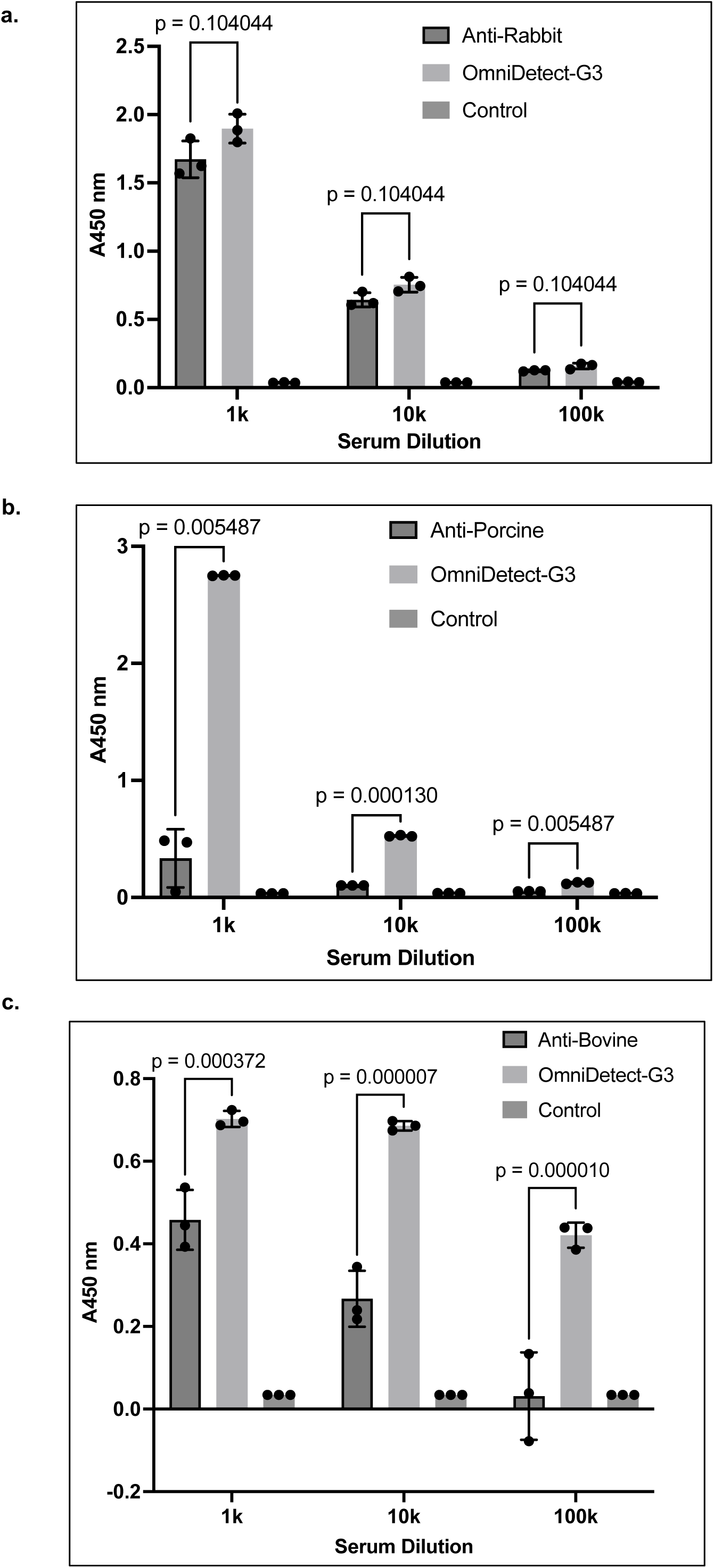
Validation of OmniDetect-G3: Detection of specific IgG in vaccinated animal serum. **a-c.** Serum from vaccinated animals containing antigen-specific IgG was collected. The corresponding purified antigen was coated on ELISA plates, and serially diluted serum was incubated with immobilized antigen. Binding was detected using either commercial HRP-conjugated secondary antibodies or OmniDetect-G3 for the comparison. Colorimetric signals were measured at 450 nm and plotted. OmniDetect-G3 vs. anti-rabbit IgG (**a**), OmniDetect-G3 vs. anti-porcine IgG (**b**), and OmniDetect-G3 vs. anti-bovine IgG (**c**). Data represent mean ± SE (n = 3). Error bars indicate standard error. P-values were calculated by Student’s t-test.

There has been a growing interest in using IgY for applications such as biopharmaceuticals and diagnostic kits^38^. Although the GB1 domain is generally thought to bind only IgG, recent studies suggested that the GB domain can weakly interact with chicken IgY. We tested whether OmniDetect-G3 can detect IgY in ELISA. Indeed, OmniDetect-G3 detected IgY at approximately 25%-40% (p<0000.1) of the signal intensity obtained with commercial HRP-conjugated anti-chicken IgY antibody (Fig. 5a).

**Fig. 5.**
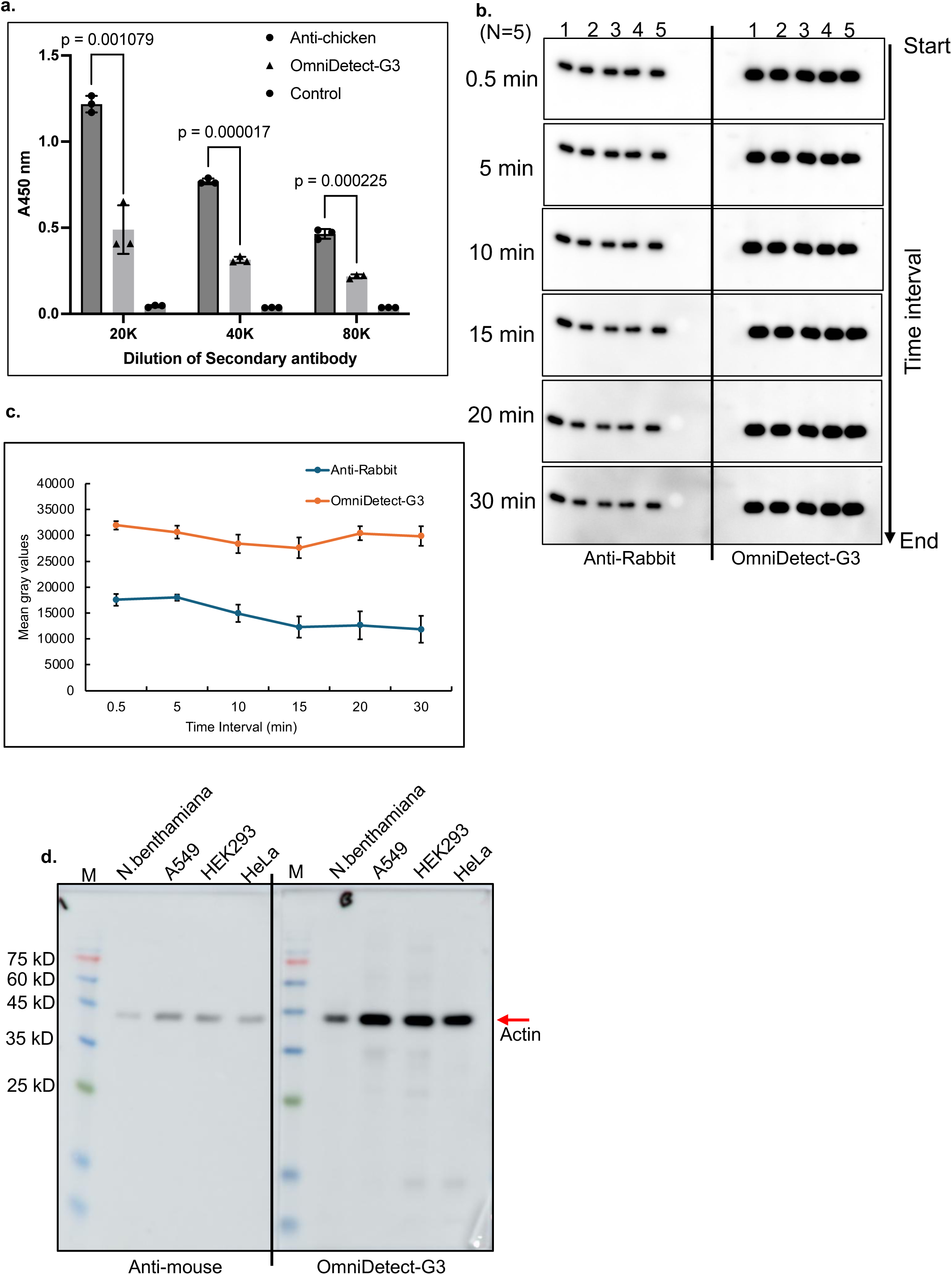
Additional validation of OmniDetect. **a.** Detection of IgY by OmniDetect. Fifty ng of highly purified His-GFP was coated on an ELISA plate and probed with primary chicken anti-His antibodies, followed by detection with serially diluted either HRP-conjugated anti-IgY antibody or OmniDetect-G3. Colorimetric signals were measured at 450nm and plotted. Data represent mean ± SE (n = 3). P-values were calculated using Student’s t-test. **b-c.** OmniDetect exhibits stable chemiluminescence over time. Fifty ng of total soluble protein from *N. bentaminana* leaves expressing His-GFP was analyzed by western blot. Band intensity was quantified using ImageJ software, and mean gray values were plotted. Data represent mean ± SE (n = 5). Error bars indicate standard error. **d.** OmniDetect shows no cross-reactivity with plant or animal cell proteins. Total soluble protein extracted from *N. benthamiana*, A549, HEK293, and HeLa cells was separated on SDS-PAGE, followed by western blotting and probed with mouse anti-actin antibody, and then incubated with either a commercial HRP-conjugated anti-mouse antibody or OmniDetect-G3. Chemiluminescence images were acquired simultaneously under a single auto-exposure setting.

### OmniDetect-G’s chemiluminescence signals persist longer and show no cross-reactivity to plant and animal total soluble proteins

During western blot analysis, we observed that the chemiluminescent signal produced by OmniDetect-G3 remained stable for an extended period of time (Fig. 5b). To compare the signal stability, we monitored the signal intensities generated by a commercial secondary antibody and OmniDetect-G3 over a 30-minute time course. Identical blots were prepared and imaged at 0.5, 5, 10, 15, 20, and 30-minute intervals using the same auto-exposure settings. For the commercial HRP-conjugated anti-rabbit IgG antibody, signal intensity was highest at 0.5 and 5 min and then gradually declined; by 30 min, the signal intensity was decreased by 33% (Fig. 5c). In contrast, the signal produced by OmniDetect-G3 remained essentially constant, showing only a 7% reduction at 30 minutes. These results suggest that the HRP moiety in the OmniDect-G3 may retain activity for a longer duration than that in the commercial secondary antibody.

We also examined the non-specific cross reactivity of OmniDetect-G3 using total soluble protein extract from the plant leaf of *N. benthamiana*, and animal cell lines–A459, HEK293, and HeLa (Fig. 5d). OmniDetect-G3 generated strong, specific signals in western blots without noticeable background cross reactivity. Although a faint smear was observed, it could be readily removed by adjusting the dilution factor of OmniDetect-G3.

## Discussion

In this study, we developed OmniDetect-G3, a plant-produced, high-sensitivity recombinant immunodetection reagent designed to replace conventional animal-derived HRP-conjugated secondary antibodies for IgG detection (Fig. 6). By fusing the GB1 immunoglobulin-binding domain with Horseradish peroxidase (HRP) and defined multimerization motifs, we systematically improved avidity and signal intensity through structural optimization. Iterative engineering—from the CTB-based pentamer (OmniDetect-G1) to the Tenascin-C–based hexamer (OmniDetect-G2) and ultimately the spatially optimized OmniDetect-G3—yielded a reagent matching or surpassing commercial HRP-conjugated secondary antibodies used in this study in western blot and ELISA assays (Figs. 1–4).

**Fig. 6.**
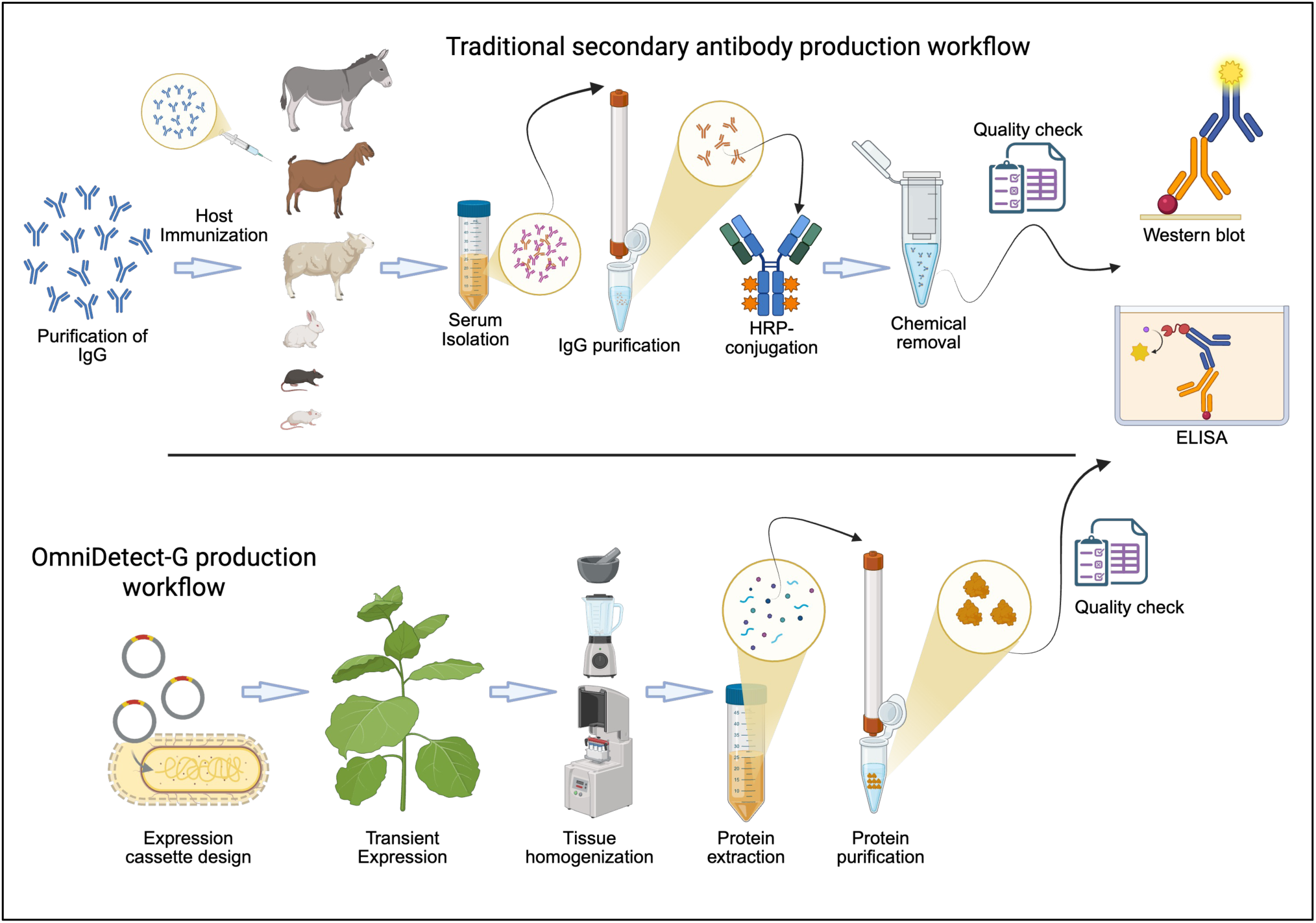
Production workflow of OmniDetect-G. Conventional secondary antibody production involves a multistep, animal-dependent process beginning with IgG antigen purification and host immunization. Achieving sufficient anti-IgG titers often takes weeks to months. After blood collection and serum extraction, IgGs must be purified. In a separate process, HRP must be purified from plant extracts. These purified IgGs and HRP are subjected to chemical conjugation, followed by further purification to remove unbound reagents. In contrast, OmniDetect-G enables a simple, rapid, and fully plant-based production workflow. The entire process-from expression to final product- can be completed in as little as two weeks. Production consists of only two steps: expression and purification, without the need for labor-intensive downstream processing beyond routine quality control (purity and activity assays). Conceptual design was created in BioRender.

The GB1 domain of *Streptococcal* protein G binds to the Fc region of IgG across diverse mammalian species but exhibits relatively weak monomeric affinity in the micromolar range^22^. Previous Fc-binding motifs – including protein A, protein L, or Z-domains-also suffer from limited affinities, restricting their utility as standalone detection reagents. To address this limitation, we employed multimerization, a principle similarly used in commercial antibody conjugates. However, unlike the non-specific chemical cross-linking used in traditional antibodies, our genetically encoded fusion strategy incorporates well-defined multimerization domains, enabling precise stoichiometry and consistent batch-to-batch performance.

OmniDetect-G1 incorporated three GB1 copies per subunit, which assembled into a pentameric complex through the CTB domain, producing a structure with 15 GB1 and 5 HRP molecules. This multimeric architecture achieved sensitivity comparable to that of commercial HRP-anti-mouse IgG and lower than anti-rabbit IgG in western blot analysis (Fig. 1i,j), while producing equal or higher ELISA signals (Fig. 1k). These findings demonstrate that avidity-driven multimerization compensates for the low intrinsic affinity of GB1, consistent with previous observation from multivalent biosensor designs.

However, OmniDetect-G1 showed lower sensitivity in western blot analysis than in ELISA, suggesting that detection efficiency depends not only on binding affinity but also on enzyme density and spatial accessibility of GB1 copies to target-bound primary antibodies^39^. To improve valency and structural flexibility, we replaced CTB with the Tenascin-C N-terminal domain (TN106), which forms a hexabrachial assembly with six extending arms. This yielded OmniDetect-G2, comprising 18 GB1 domains and 6 HRP molecules per complex (Fig. 2b). The enhanced geometric arrangement likely reduced steric hindrance and mitigated potential HRP self-quenching, resulting in ELISA signals up to 101% stronger than those generated by HRP-anti-rabbit IgG (Fig. 2k). Notably, the plant-derived HRP retained N- N-glycosylation, providing improved folding and stability^29^ compared with microbial HRP variants (HRP4m, HRP6m)—highlighting a key advantage of plant molecular farming^40^.

Building on these findings, we optimized GB1 positioning by placing domains at both termini of TN106—one at N-terminus and two at C-terminus—resulting in creation of OmniDetect-G3 (Fig. 3a). This configuration improved GB1 accessibility and enhanced cooperative avidity, producing markedly higher sensitivity: 165–381% greater in western blot analysis and 30%–1,145% higher in multi-species ELISA compared to commercial antibodies (Fig. 3h–4c). These results suggest that OmniDetect-G3 outperforms previously reported HRP–protein G or HRP–ABD fusions.

Time-dependent durability testing showed that OmniDetect-G3 retained over 93% of its signal after 30 minutes of substrate incubation, whereas conventional HRP-conjugated secondary antibodies exhibited a 33% signal loss over the same period (Fig. 5c). The radially distributed arrangement of HRP moieties within the hexabrachial scaffold likely reduces localized oxidative stress, enhancing enzyme longevity.

A key advantage of OmniDetect-G3 is its host-independent recognition. Unlike species-specific secondary antibodies (e.g., goat anti-rabbit or donkey anti-mouse), OmniDetect-G3 effectively detects IgGs from rabbit, mouse, pig, and cattle, and also partially cross-reacts with chicken IgY (∼25% of standard anti-IgY signal) (Fig. 5a). Although GB1 alone binds IgY only weakly, multimerization significantly improves its apparent affinity, suggesting that further engineering could enhance IgY detection.

Compared with recombinant HRP-Fc-binding constructs expressed in *E. coli* or *Pichia pastoris*, the OmniDetect-G3 platform benefits from hexameric architecture, higher HRP valency (six per complex), and plant-specific post-translational modifications that preserve catalytic activity. These features collectively support low-cost, sustainable, animal-free production-aligning well with the increasing emphasis on ethical biomanufacturing.

Collectively, OmniDetect-G3 represents a versatile and sustainable alternative to animal-derived secondary antibodies. Beyond western blotting and ELISA, future directions include fusion with alternative reporters (e.g., alkaline phosphatase, luciferase) to enable multimodal detection. Integration into lateral-flow or electrochemical biosensor systems could further expand its diagnostic applications. By combining high sensitivity, enhanced stability, and ethical and animal-free production, OmniDetect-G3 establishes a next-generation framework for universal immunoassay reagents that balance performance, reproducibility, and sustainability.

## Materials and Methods

### Construct design

The nucleotide sequences of *CTB* (WP_338286491.1), *TN106* (encoding 34-139 aa; Uniprot ID: P10039), and the three *HRP* variants (encoding 31-338 aa; Uniprot ID: P00433) -WT, 6m, and 4m-were chemically synthesized by Gene Universal. A 6x His tag and an HDEL motif were added at the 3’ of HRP to enable Ni^2+^-NTA affinity purification and ER retention, respectively. For subcloning of *HRP*, BamHI and XhoI restriction endonuclease sites were added at the 5’ and 3’ ends, respectively. DNA fragments containing zein leader sequence (*Z*), *CTB*, and the three copies of *GB1* (*GBx3*), as well as fragments containing ZTN106 and GBx3, were also synthesized with XbaI and BamHI restriction endonuclease sites incorporated at the 5’ and 3’ends. The *BamHI-HRP-6xHis-HDEL-XhoI* fragment was digested with BamHI and XhoI and ligated to the pFM’M vector that had been digested with the same restriction endonucleases, resulting in the construct *HRP-6xHis-HDEL*.

The pUC plasmids carrying XbaI-Z-CTB-GBx3-BamH1 or XbaI-Z-TN106-GBx3-BamH1 were digested with XbaI and BamHI, and the resulting DNA fragments were ligated to the pHRP-6xHis-HDEL vector that had been digested with the same restriction endonucleases, yielding pZCTB-GBx3-HRP-6xHis-HDEL and pZCTB-GBx3-HRP-6xHis-HDEL, designated as pOmniDetect-G1 and pOmniDetect-G2, respectively. To generate pOmniDetect-G2-4m and pOmniDetect-G2-6m, the wild-type HRP region was replaced by chemically synthesized *HRP-4m* or *HRP-6m* using BamHI and XhoI restriction endonucleases. For the construction of pOmniDetect-G2-C, a full-length fragment containing *XbaI-Z-TN106-HRP-GBx3-6xHis-HDEL-XhoI* was chemically synthesized, digested with XbaI and XhoI, and ligated to pOmniDetect-G2 that had been digested with the same restriction endonucleases. To generate OmniDetect-G3, a DNA fragment encoding XabI-Z-GB1-TN106-GBx2-BamH1 was synthesized, digested with XbaI and BamH1, and then ligated into pOmniDetect-G2, digested with the same restriction endonuclease to generate pZ-GB-TN106-GBx2. Subsequently, the wild-type *HRP-6xHis-HDEL* fragment flanked by BamH1 and XhoI at 5’ and 3’ ends, respectively, was synthesized, digested with BamH1 and XhoI, and ligated into pZ-GB-TN106-GBx2 that had been digested with BamHI and XhoI to generate pZ-GB-TN106-GBx2-HRP-6xHis-HDEL, designated as OmniDetect-G3.

### Plant Growth Conditions

Wild-type *N. benthamiana* plants were cultivated in a controlled greenhouse environment. The temperature was maintained at 24°C with the relative humidity of 40–65%. A long-day photoperiod was used, consisting of 14 hours of light and 10 hours of darkness (light intensity, 130–150 μE/m2), for a duration of 5 to 7 weeks. For agro-infiltration, 5 to 7-week-old plants were used.

### Agroinfiltration, protein expression, and analysis

Transient expression of OmniDetect variants was achieved in *N. benthamiana* leaves. All expression vectors were introduced into *Agrobacterium tumefaciens* strain EHA105 by electroporation. A single colony was used to inoculate 5 mL of LB medium (LPS Solution, Cat. LBL-05) and cultured overnight. The overnight culture was then added to 50 ml of LB medium containing appropriate antibiotics. After pelleting the cells by centrifugation at 3,500 × g for 10 min, the pellet was resuspended in infiltration buffer (10 mM MES, 10 mM MgSO_4_, 200 μM acetosyringone, pH 5.6) to an optical density of 0.6-0.7 at 600 nm. The cell suspension was incubated at room temperature for a period of 2 to 4 h before infiltration. The abaxial side of leaves from 5–7-week-old plants was infiltrated using a 1 ml needleless syringe or via vacuum infiltration. Infiltrated plants were returned to the greenhouse and incubated for an additional 3 to 7 days.

Leaf tissues were harvested, ground into a fine powder using a bead beater at 20 Hz for 1 min under liquid nitrogen or with a mortar and pestle for large sample quantities. The powdered tissue was mixed with 5 volumes (w/v) of protein extraction buffer (50 mM Tris-HCl, pH 7.4, 300 mM NaCl, 0.1% [v/v] Tween-20, and 0.5X protease inhibitor cocktail). Total soluble protein was obtained after centrifugation at 16,000 × g for 15 min, repeated twice. Protein concentration was determined using the Bradford protein assay (Bio-Rad, Hercules, CA, United States).

### SDS-PAGE and western blot analysis

Proteins were separated by 5 or 12% SDS-PAGE. Gels were either stained with 0.25% CBB R-250 solution (AMRESCO, catalog number: 6104-59-2) containing 45% methanol and 10% glacial acetic acid or subjected to western blot analysis. For Western blot analysis, membranes were blocked in a solution of 3% fat-free skim milk prepared in TBST buffer (20 mM Tris-Cl, pH 7.5, 150 mM NaCl, 0.1% Tween-20) for 1 h at room temperature. Membranes were then incubated overnight at 4 °C with one of the following primary antibodies diluted 1:1,000 in TBST containing 3% non-fat dry milk: mouse anti-His (Novus, AD1.1.10), rabbit anti-GFP (GenScript, Cat# A01388, USA), mouse anti-GFP (Living Colors JL-8, Cat# 632381, USA), or chicken anti-His (abcam, Cat# ab9107, United Kingdom). After three washes with TBST, membranes were incubated for 1 h at room temperature with appropriate secondary antibodies diluted at 1:10,000 or with OmniDetect diluted at 1:50,000-1:80,000 in TBST without skimmed milk. Immunoblot signals were visualized using Amersham Imager 680 (GE Healthcare, Chicago, IL, United States).

### Protein purification using Ni^2+^-NTA affinity chromatography

Total soluble extract from 30 g of infiltrated leaves was prepared as explained above. Four ml of Ni^2+^-NTA resin (Qiagen, Cat.#30230, Germany) were mixed with the extract and incubated at 4°C for 2 h with gentle agitation to allow protein binding. The mixture was then slowly passed through a glass column. The unbound fraction was collected, and a 2-ml aliquot was stored at −80°C for further analysis. The Ni^2+^-NTA-bound OmniDetect protein was washed with 100 ml of washing buffer (50 mM Tris-Cl, pH 7.4, 300 mM NaCl) containing 30 mM imidazole. A 2-ml of the wash fraction was stored at −80°C for analysis. Before elution, resin-bound protein was washed with 10-20 ml of Tris-Cl (pH 6.0) to remove residual salts. OmniDetect proteins were eluted using a stepwise imidazole gradient (50 mM, 100 mM, and 250 mM) prepared in Tris-Cl (pH 6.0) buffer. The eluted fractions were pooled and concentrated using a 30 kDa Amicon Ultra-15 centrifugal filter (Merk Millipore, Cat.# UFC9030, USA).

### Serum samples from vaccinated animals

BioApplications Inc., Pohang, Gyeongsangbuk-do, Korea, kindly provided serum samples containing anti-porcin-CSF IgG and anti-bovine tuberculosis Ag85A. These sera samples were collected from vaccinated animals in the field, not from experimental animals. Therefore, no permission is required for animal experiments. Anti-GFP IgG was collected from rabbit sera used in the experiments at Gyeongsan National University, Korea (Approval number: GNU-25024-B0016).

### ELISA assay

Fifty ng of purified His-GFP, porcin-CSF antigen, and bovine tuberculosis Ag85A proteins were coated onto a 96-well microtiter plate and incubated overnight at 4°C. After blocking with 200 µL of 3% BSA in TBST buffer for 1 h at room temperature, the corresponding serum (diluted 1:1000-1:100,00) from vaccinated animal or primary antibody (diluted 1:5,000) was added and incubated on a microtiter plate shaker for 2 h at 37°C. Following five washes with 250 µL TBST buffer, HRP-coupled secondary antibodies-anti-mouse, anti-rabbit, anti-porcine, anti-bovine, anti-IgY – diluted at 1:10,000-1:640,000, or OmniDetect diluted 1:10,000-1:640,000, were added and incubated for 1 h at 37°C. After five additional washes with TBST buffer, 50 µL of TMB (3,3ʹ,5,5ʹ-tetramethylbenzidine) substrate (Thermo Fisher, Cat# 34025) was added and incubated for 30 s to 5 min, followed by addition of 50 µL 0.18 M H_2_SO_4_ to stop the reaction. Absorbance was measured at 450 nm using a plate reader.

### Band intensity calculation and statistical analysis

Mean gray values of SDS-PAGE gels and western blot membranes were quantified using ImageJ software. Statistical analyses and p-values were generated using GraphPad Prism version 10.0.0 (GraphPad Software, Boston, Massachusetts, USA).

### Conceptual protein structure design of OmniDetect-G

The conceptual 3D models of OmniDetect were generated using a licensed version of Biorender. Protein structures were imported from the PDB database using Biorender’s built-in import function. The following PDB IDs were used: CTB: 1PZJ, GB1: 1GB1, HRP: 1HCH. The final conceptual illustrations represent schematic models, and the actual 3D crystal structure of OmniDetect may differ from these representations.

## Acknowledgements

This work was supported by grants (RS-2023-00235511, RS-2025-16066359, and RS-2025-02634433) from the National Research Foundation of Korea, Ministry of Science and ICT, Korea.

## Author contribution

Hwang I. and Soni A.P. contributed to conceptualizing this study and wrote the manuscript; Soni A.P. and Kumari M. made DNA constructs and contributed significantly to performing the experiments and analyzing the results. T.M. and Y.H. performed some tests of OmniDetect-G3 with cell materials. S.W. established protein purification conditions. All authors read and approve the manuscript.

