## Supplementary material for "OmniDetect: a plant-produced universal IgG binder replacing secondary antibodies in immunoassays": Supplementary Figures.pdf

**Supplementary Fig. 1**

a.

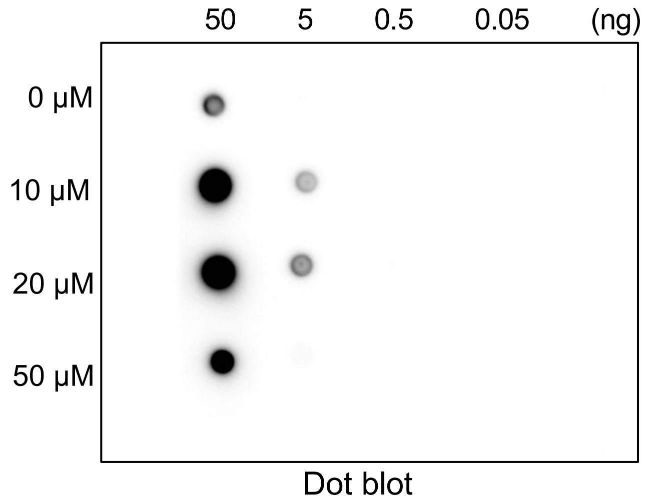


b.

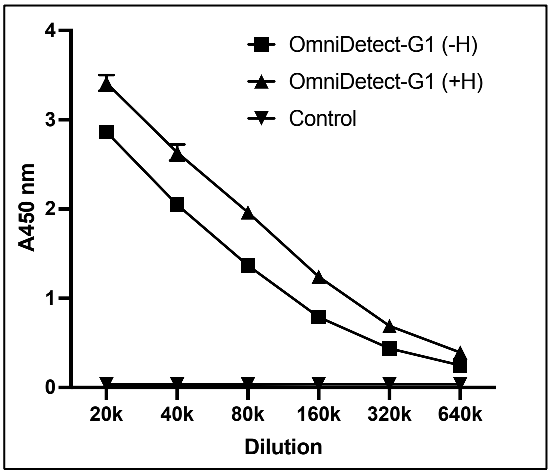


**Supplementary Fig. 1:** Hemin treatment of OmniDetect-G1. **a.** Optimization of hemin concentration. Purified OmniDetect-G1 (1 mg/ml) was incubated with various concentrations of hemin. After incubation, buffer exchange was performed to remove the residual hemin, and samples were analyzed by dot-blot. Hemin at 10-20 μM (final concentration) produced the strongest signals upon ECL development, whereas signal intensity decreased at 50 μM. **b.** ELISA before and after hemin incorporation. Fifty ng of His-GFP was coated on an ELISA plate and probed with rabbit-anti-GFP IgG, followed by serially diluted OmniDetect-G1 with (+H) or without (-H) hemin treatment. Hemin incorporation significantly increased ELISA signal intensity across all dilutions ($p < 0.0000001$; at 40,000 dilution). Data represent mean ± SE (n = 3). For control, standard deviation bars are not visible due to extremely low values.

**Supplementary Fig. 2**

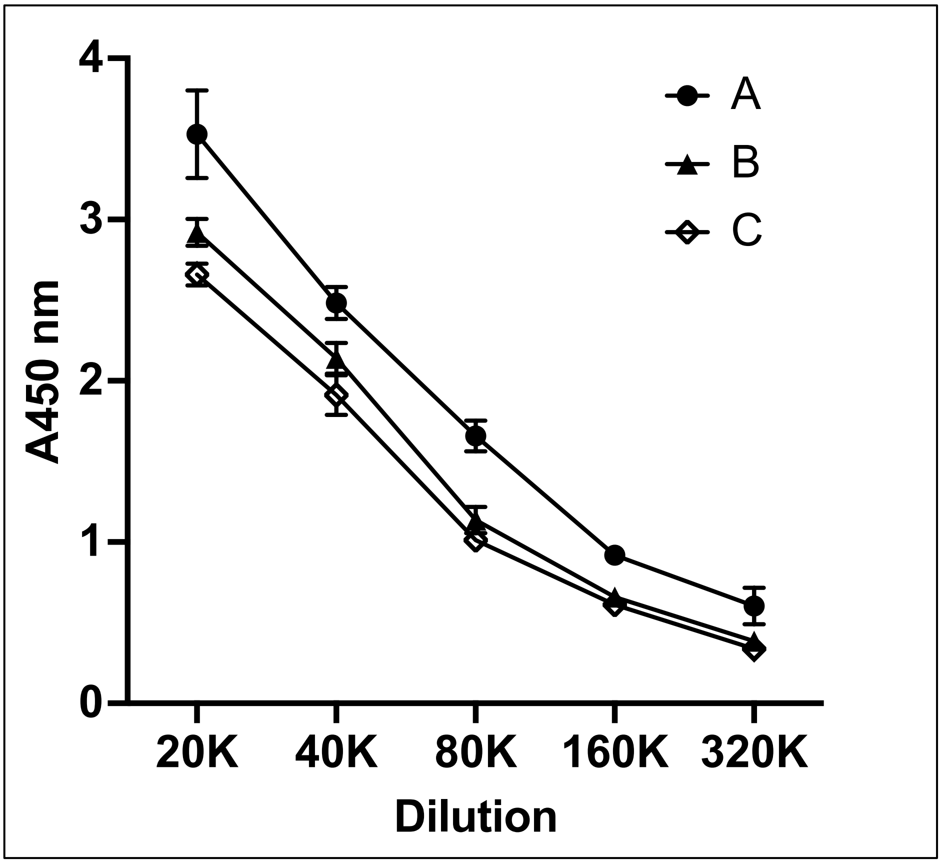


**Supplementary Fig. 2. Selection of the best commercial secondary antibody**.

Fifty ng of purified His-GFP was coated on an ELISA plate and probed with Rabbit-anti-GFP IgG. HRP-conjugated anti-rabbit antibodies from three different companies (named here as A, B, and C) were serially diluted and incubated with the samples. Colorimetric signals were measured at 450nm and plotted. Data represent mean ± SE (n = 3).

**Supplementary Fig. 3**

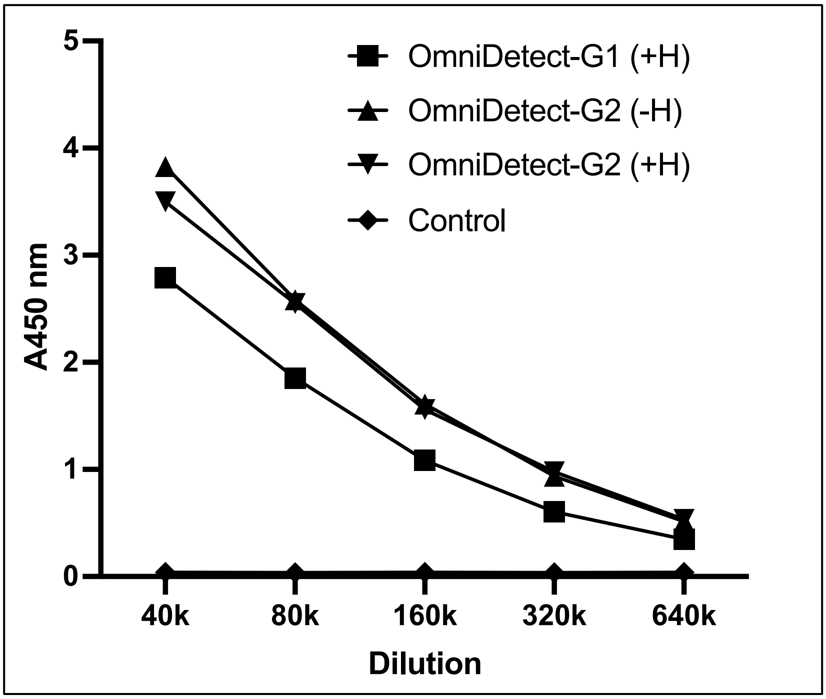


**Supplementary Fig. 3. Comparative ELISA analysis of OmniDetect-G1 and OmniDetect-G2.** Hemin-treated OmniDetect-G1 (+H; used as controls) was compared with OmniDetect-G2 with or without hemin treatment in ELISA. OmniDetect-G2 showed no significant difference in the activity regardless of hemin incorporation, whereas OmniDetect-G2 exhibited significantly higher activity compared to OmniDetect-G1 (+H) ($p < 0.001$). Data represent mean ± SE ($n = 3$). For control, standard deviation error bars are not visible due to extremely low values. P-values were calculated using a two-tailed Student's *t*-test.

### Supplementary Fig. 4

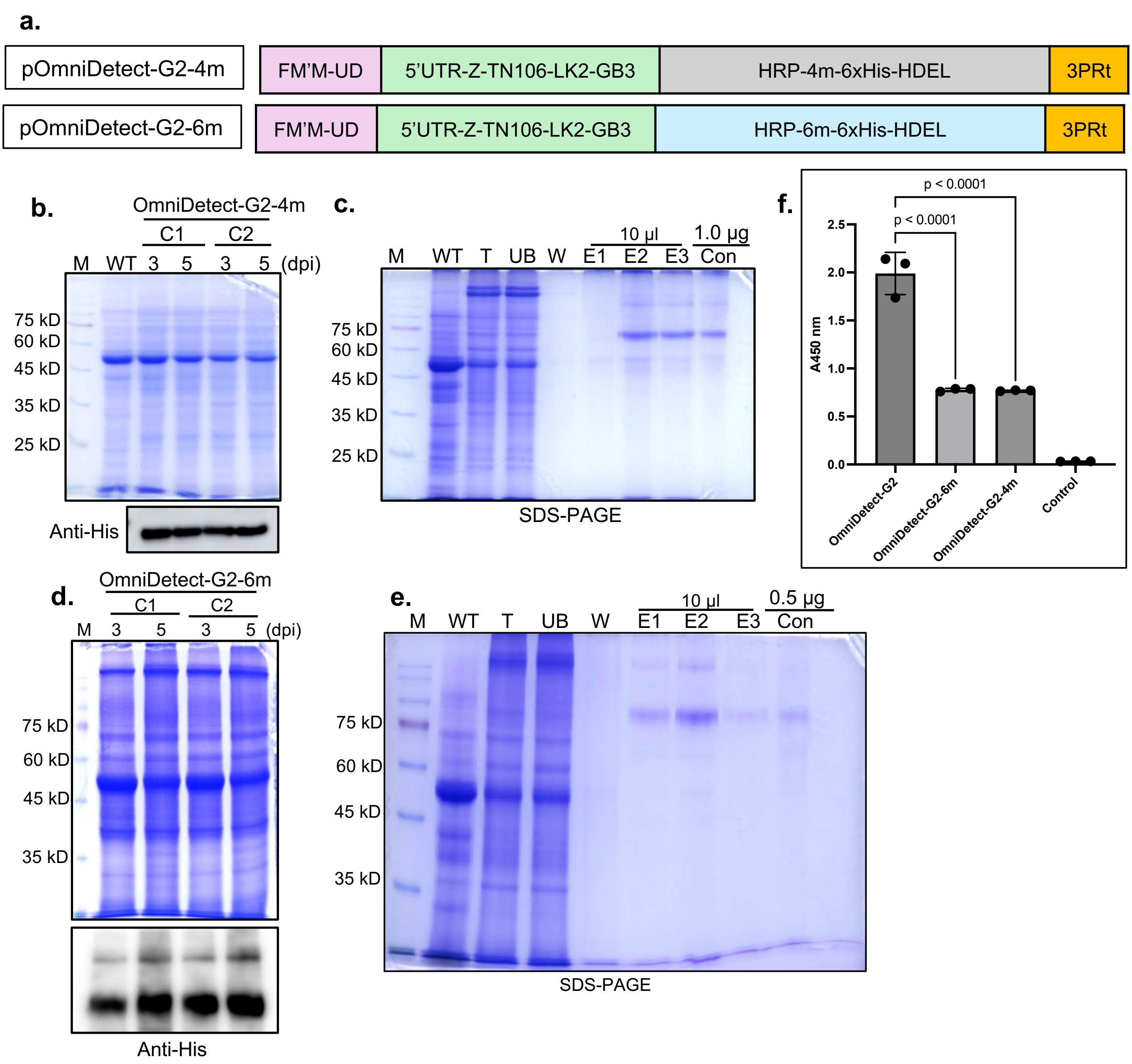


**Supplementary Fig. 4: Characterization of mutant OmniDetect-G2 variants. a.** Schematic representation of construct designs. **b,c.** Protein expression in two independent colonies, and purification of OmniDetect-G2-4m. **d,e.** Expression of OmniDetect-G2-6m and purification using $Ni^{2+}$-NTA affinity chromatography. **f.** Comparative ELISA analysis of OmniDetect-G2, OmniDetect-G2-4m, and OmniDetect-G2-6m. Data represent mean ± SE (n = 3). For control, the standard deviation error bars are not visible due to extremely low values. P-value was calculated using one-way ANOVA. M, protein marker; WT, wild-type protein extract; C, colony; T, total soluble protein; UB, unbound fraction; W, washing fraction; E1, E2, and E3, eluted fractions with 50, 100, and 250 mM imidazole, respectively; Con, concentrated protein using Centricon.

**Supplementary Fig. 5**

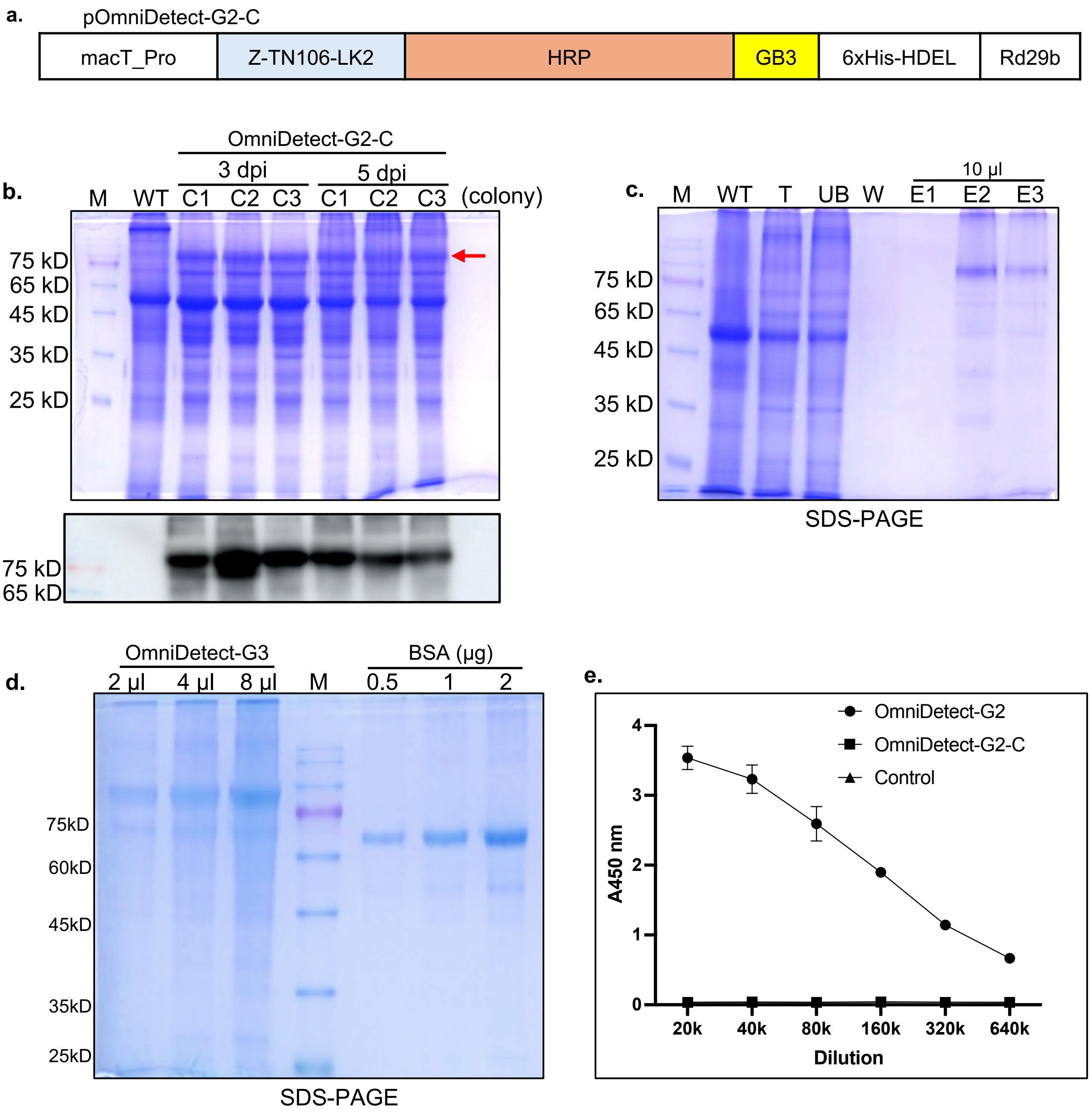


**Supplementary Fig. 5: Characterization of OmniDetect-G2-C. a.** Schematic representation of construct design; **b-d.** Protein expression in three independent colonies, purification, and quantification of OmniDetect-G2-C. **e.** Comparative ELISA analysis of OmniDetect-G2 and OmniDetect-G2-C. Data represent mean ± SE (n = 3). For control and G2-C, the standard deviation error bars are not visible due to extremely low values.

M, protein marker; WT, wild-type protein extract; C, colony; T, total soluble protein; UB, unbound fraction; W, washing fraction; E, elution fraction; Con, concentrated protein using Centricon.
